# SCMarker: ab initio marker selection for single cell transcriptome profiling

**DOI:** 10.1101/356634

**Authors:** Fang Wang, Shaoheng Liang, Tapsi Kumar, Nicholas Navin, Ken Chen

## Abstract

Single-cell RNA-sequencing data generated by a variety of technologies, such as Drop-seq and SMART-seq, can reveal simultaneously the mRNA transcript levels of thousands of genes in thousands of cells. It is often important to identify informative genes or cell-type-discriminative markers to reduce dimensionality and achieve informative cell typing results. We present an *ab initio* method that performs unsupervised marker selection by identifying genes that have subpopulation-discriminative expression levels and are co- or mutually-exclusively expressed with other genes. Consistent improvements in cell-type classification and biologically meaningful marker selection are achieved by applying SCMarker on various datasets in multiple tissue types, followed by a variety of clustering algorithms. The source code of SCMarker is publicly available at https://github.com/KChen-lab/SCMarker.

**Author Summary:** Single cell RNA-sequencing technology simultaneously provides the mRNA transcript levels of thousands of genes in thousands of cells. A frequent requirement of single cell expression analysis is the identification of markers which may explain complex cellular states or tissue composition. We propose a new marker selection strategy (SCMarker) to accurately delineate cell types in single cell RNA-sequencing data by identifying genes that have bi/multi-modally distributed expression levels and are co- or mutually-exclusively expressed with some other genes. Our method can determine the cell-type-discriminative markers without referencing to any known transcriptomic profiles or cell ontologies, and consistently achieves accurate cell-type-discriminative marker identification in a variety of scRNA-seq datasets.

## Introduction

Current single-cell RNA-sequencing (scRNA-seq) data generated by a variety of technologies such as Drop-seq and SMART-seq, can reveal simultaneously the mRNA transcript levels of thousands of genes in thousands of cells [1–3]. However, the increased dimensionality makes it challenging to delineate cell types, due to complex and often undefined associations between individual genes and cell-types [4,5]. It is well accepted that genes are not equally informative in delineating cell types [6,7]. Certain genes are only expressed in certain cell types, but not others [8]. Moreover, the expression levels of certain genes cannot be robustly measured (e.g., zero inflated), due to technological bias [9–11]. Thus, it has become a common practice to retain only highly expressed or highly variable genes for cell populational analysis [12–15]. Several scRNA-seq data clustering packages (**S1 Table**) perform marker selection through dimensionality reduction techniques such as principal component analysis and tSNE [16], which are equivalent to identifying the set of highly variable genes. Unfortunately, the biological implications and the technical optimality of these gene selection strategies retain unclear, despite their wide use in cell-type clustering.

Here, we propose an *ab initio* method, named SCMarker, which applies information-theoretic principles to determine the optimal gene subsets for cell-type identification, without referencing to any known transcriptomic profiles or cell ontologies. The central idea of our method is to select genes that are individually discriminative across underlying cell types, based on a mixture distribution model, and are co- or mutually exclusively expressed with some other genes, due to cell-type specific functional constraints. Although the techniques of applying a mixture distribution model for a set of continuous data points have been widely used in clustering analysis of gene expressions, it is unclear whether this approach can benefit this problem context [17,18]. In particular, because single-cell gene expression measurements have vast dimensions (>20,000 genes), are highly noisy (e.g., zero-inflated, drop-off errors), and are generated by technologies of varied properties [19,20]. For example, SMART-seq is aimed at sequencing the entire RNA transcript, while Drop-seq only the 3’ end using unique molecular indices (UMI) to track individual transcript [2,21,22]. Part of our investigation here is to examine whether the previously applied data analytical techniques can be reapplied in the single-cell data-type that have different properties and population structures. Our main goal is to identify not only cell-types, but also biologically meaningful cell-type markers from scRNA-seq data at accuracies higher than results derived using canonical gene selection strategies.

## Materials and Methods

### Discriminativeness of gene expressions for subpopulation clustering

By definition, cell-type-discriminative markers (CTDMs) should have distinctive expression levels across cell subpopulations. Therefore, in a dataset with mixed cell subpopulations, the expression level of a CTDM should follow a bi- or multi-modal, instead of a unimodal distribution (**Fig. 1A and B**) [23–25]. Following this assumption, we quantify the degree of modality based on the probability density distribution (*f*) of each gene expression using a Gaussian kernel function, instead of a mixture model which requires knowing the number of mixture components:

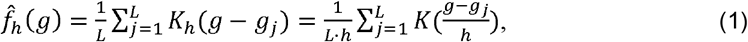

where *g*_*j*_ is the expression level of a gene in cell *j*; *h*> 0 is a smoothing parameter called bandwidth, which is estimated by the average level of gene expression in cell population and can be leveraged to alleviate biases introduced by sequencing depths; and *L* is the number of cells. *K*_*h*_ (*x*) is a scaled kernel function defined as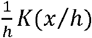, where *K*(*x*) is a standard Gaussian density function. For each gene, we count the number (*T*) of peaks in the estimated probability density function 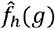. A peak is found at the density value *c*, if there exists a 2 times *h* long interval *I* centred at *c* such that 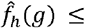 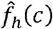 for all *g* in *I*. A gene expression level follows a multi-modal distribution, if it has multiple (*T* ≥ 2) local maximum probability density values. Only genes with multimodal probability density distributions are considered as markers.

**Fig. 1.**
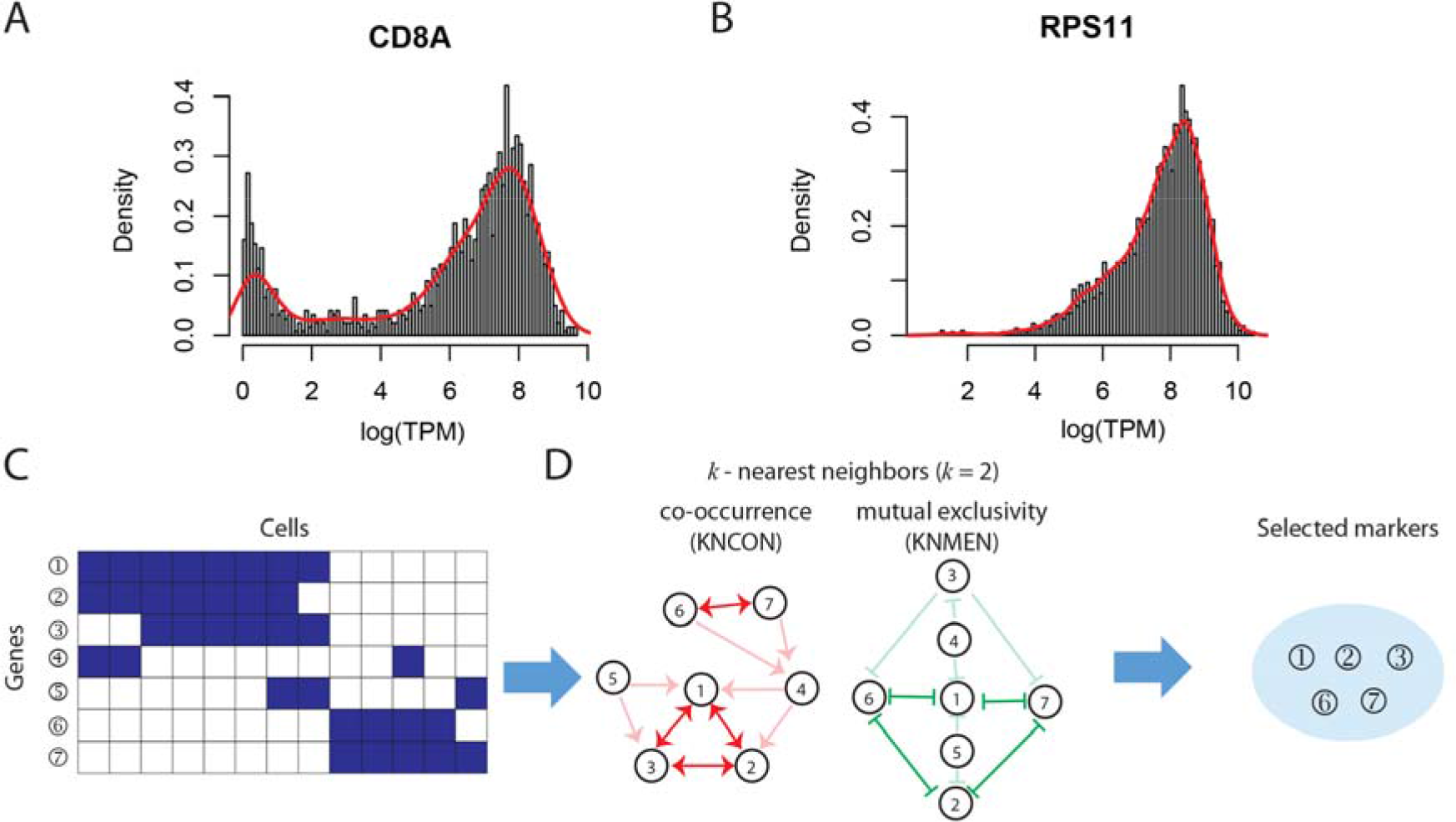
Illustration of SCMarker. Plotted as examples are (**A**) a bimodally distributed gene expression and (**B**) a unimodally distributed gene expression. From a binarized gene-cell expression matrix (**C**), a *k*-nearest co-occurrence neighbour (KNCON) graph and a *k*-nearest mutually exclusive neighbour (KNMEN) graph are constructed (**D**), based on which co- or mutually exclusively expressing gene pairs (CMEGPs) in the KNCON (node 1, 2 and 3, node 6 and 7, connected by red double arrows) and in the KNMEM (node 1, 2, 6 and 7, connected by green double arrows) can be identified. Marker genes (node 1, 2, 3, 6 and 7) are subsequently selected based on the CMEGPs.

### Co- or mutually exclusively expressing gene pairs (CMEGPs)

CTDMs are often co- or mutually exclusively expressed, due to modularized regulatory interactions specific to cell types [26]. Consequently, identifying these CMEGPs, will help identify CTDMs. Because scRNA-seq data are often sparse with limited sequencing depth [27], binarization of the counts would help mitigate technical artifacts and improve robustness over different sequencing platforms (e.g., whole transcript vs 3’ sequencing protocols). To identify CMEGPs, we only consider genes with multimodal distribution and discretize a gene-cell expression matrix with *N* genes and *L* cells into an *N* × *L* binary matrix *X* ∈ {0, 1}^*N*×*L*^, with *x*_*ij*_ = 1 designating an expressed gene *i* in cell *j*, if the expression level is above the average and *x*_*ij*_ = 0 otherwise (**Fig. 1C**). For gene *i*, *x*_*i*_ = (*x*_*i1*_, *x*_*i2*_, …, *x*_*iL*_) is a binary string. We can calculate a co-occurrence matrix (*S*) that measures the pair-wise co-occurrence between all the gene pairs,

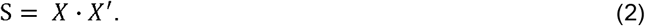

S can also be represented as a directed graph (**Fig 1D**), in which a node denotes a gene and an edge from gene A to gene B represents that B co-occurs with gene A in at least *n* cells. Among the connected nodes, the *k* genes that co-occurred with A in *k* largest sets of cells are termed the *k*-nearest co-occurrence neighbours (KNCONs).

In addition, we calculate a mutually exclusive matrix (*M*) that measures the pair-wise mutual exclusivity between all the gene pairs through equation (3),

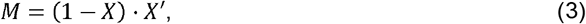

where *M* represents a directed *k*-nearest mutually exclusive neighbour (KNMEN) graph (**Fig. 1D**). Similar to KNCON, but in opposite ways, the KNMENs of a gene A are the *k* genes that occur mutually exclusively with A in *k* largest sets of cells.

Under these definitions, an CMEGP is identified as two genes bi-directionally connected in the KNCEN or the KNMEN graph. We selected as markers the genes that belong to at least one CMEGP, because these genes are more likely associated with cell-type specific functions than those that do not have any CMEGP (likely due to random, non-function-related fluctuation). This concept has been previously examined in the RNA microarray data analysis, but has not been successfully applied in the context of single-cell RNA-seq data analysis, due to vastly different properties between the technologies [17,18].

## Results

### Comparison with other marker selection strategies

We applied SCMarker to the scRNA-seq data obtained from 1) 19 melanoma patients, which include 4,645 cells; and 2) 18 head and neck cancer patients, which include 5,902 cells sequenced by the SMART-seq2 platform [28,29]. In the original studies, each cell in the sets was labelled as a malignant or non-malignant cell through copy number analysis. The expression levels of known marker genes were used to further classify the non-malignant cells, such as T cells, B/plasma cells, macrophages, dendritic cells, mast cells, endothelial cells, fibroblasts, and myocytes. We found that most known marker genes (96%) demonstrated bi/multi-modal distributions across cells (**S1 Fig.**). Overall, around 6% of genes with bi/multi-modal distributions are identified as marker genes, among which half are the known marker genes.

For a fair comparison, we assessed SCMarker results with those obtained under two canonical strategies: selecting genes with A) the highest average expression levels and B) the highest variance across cells. In our experiments, the highest variable genes were determined using Seurat [12]. We used five clustering methods: k-means, Clara, hierarchical clustering, DBSCAN and Seurat to cluster single cells based on the selected markers [30,31]. Same numbers of clusters are specified for DBSCAN, k-means, Clara, and hierarchical clustering. The adjusted rand index (ARI), which measures the similarity of two sets of clustering results, was used to quantify the consistency between the clustering results and the known cell labels [32]. Compared to marker sets A and B, selected by the canonical strategies, the marker set selected by SCMarker (equal numbers of markers) resulted in a higher ARI with fairly evident margins (Fig. 2). The conclusion appeared to be robust over a range of *k* and *n* parameters and were unaffected by using different clustering methods (S2 **Fig.**).

**Fig. 2.**
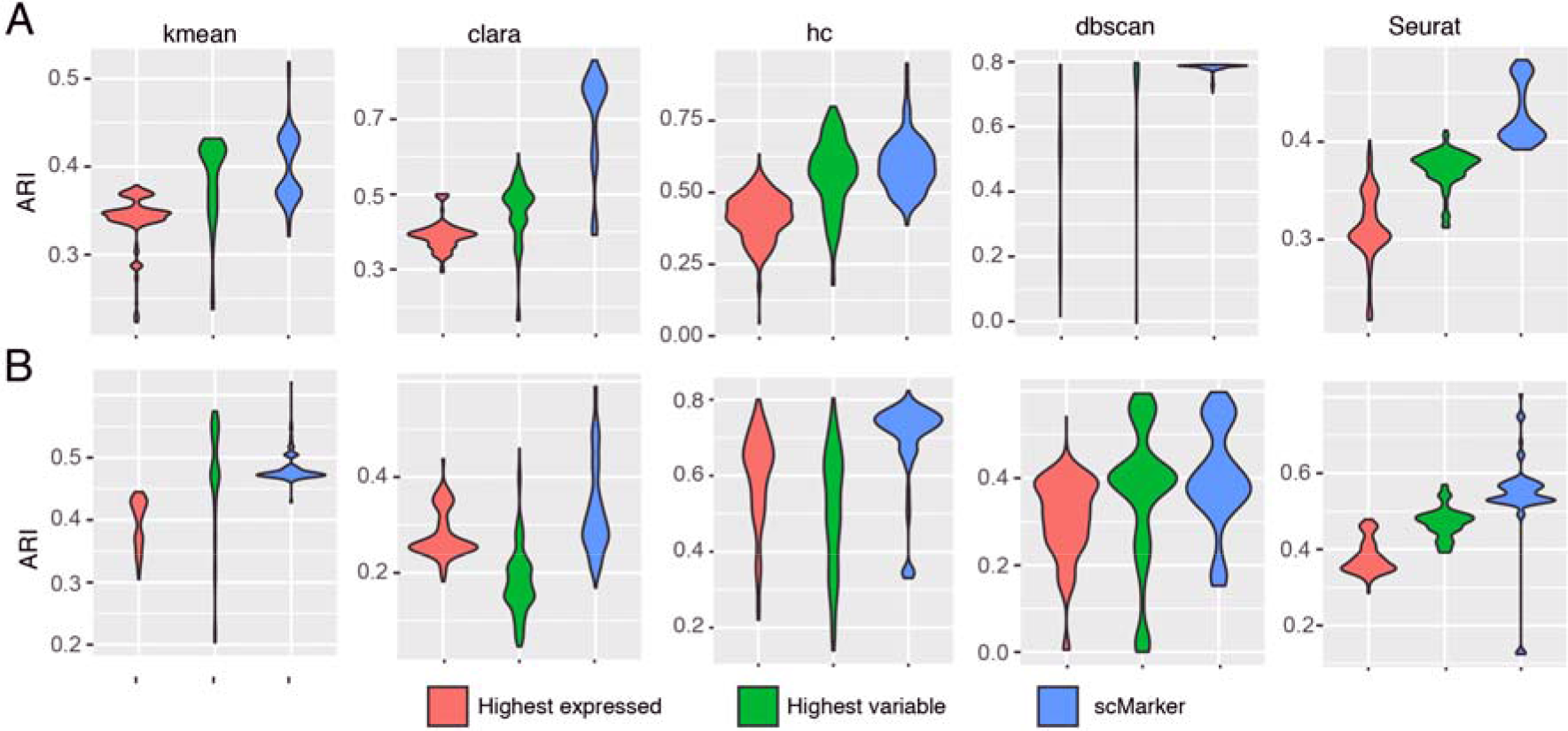
Comparison of 3 marker selection methods for cell-type identification. Accuracy of cell-type identification (in terms of adjusted rand index) are compared across 3 marker sets selected respectively by SCMarker, the highest expressed and the highest variable gene approaches, using two scRNA-seq datasets from (A) melanoma and (B) head-and-neck cancer samples by 5 clustering algorithms: k-means, Clara, hierarchical clustering (hc), DBSCAN, and Seurat.

These experiments indicated that setting *k* between 100 and 300 resulted in the most accurate cell type identification results irrespectively to *n* (**S3 Fig.**). Hence, we select *k* = 300 and *n* = 30 as the default parameters for applying SCMarker.

We obtained 902 markers from the melanoma data and more distinguishable cell types using SCMarker than using the canonical strategies (**Fig. 3A** to **C**). Better performance of SCMarker was also obtained in analysing the head and neck cancer data (**S4 Fig.**). Moreover, the genes selected by SCMarker had substantially higher degrees of overlap with the known cell-type markers reported in the original publications than the sets returned by other approaches (the same number of 902 top scoring genes were selected for fair comparison), including the “FindMarker” approach in Seurat (**Fig. 3D to G**). Notably, SCMarker selected significantly more immune cell surface markers specific to T cytotoxic, T helper, B lymphocyte, and macrophage cells that are likely present in the tumour microenvironment, as indicated by gene set enrichment analysis (**S5 Fig.**) [33].

**Fig. 3.**
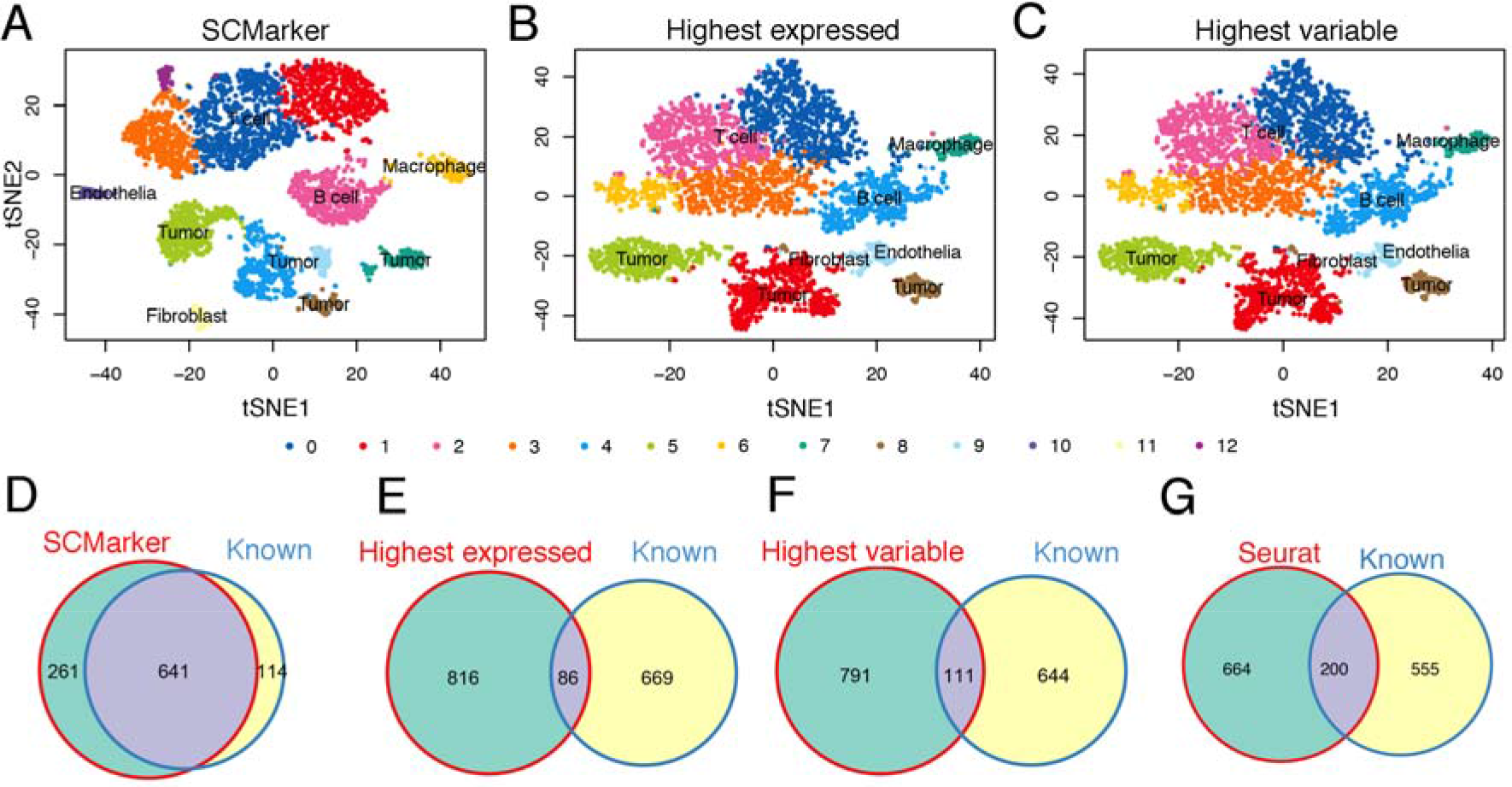
Results on the melanoma data. Plotted in tSNE space are 4,645 melanoma cells with markers selected respectively by (**A**) SCMarker, (**B**) the highest expressed and (**C**) the highest variable genes. Also plotted are the Venn diagrams between the known cell-type markers and the marker sets determined respectively by (**D**) SCMarker, (**E**) the highest expressed, (**F**) the highest variable genes and (G) Seurat FindMarker in the melanoma data.

### Application of SCMarker to 3’ UMI count data

To avoid introducing biases due to potential overfitting and assess the utility of our approaches on other platforms, we further assessed the utility of SCMarker in analysing the 3’ UMI count scRNA-seq data generated by the droplet platforms.

We first analysed a set of 5,602 cells from the cerebellar hemisphere of normal brain tissues generated by Drop-seq [34]. SCMarker selected 699 genes as CTDMs, which differentially expressed across cell subpopulations under the default parameters. Alternatively, the default mode of Seurat led to the selection of 6,111 highest variable genes (HVGs). For comparison, we selected 699 highest expressed genes (HEGs). Although SCMarker selected less markers than Seurat, the clustering result showed a clearer separation than that based on the Seurat HVGs and on the HEGs (**Fig. 4A** to **C**). In particular, SCMarker successfully delineated Purkinje neurons into purk1 (cluster4, **Fig. 4A**) and purk2 (cluster7, **Fig. 4A**) and recapitulated the differential levels of *SORCS3* between two clusters (**Fig. 4D**), which are consistent with the results in the original paper. In contrast, although the Purkinje neurons were clustered into four groups by Seurat (**Fig. 4B**), purk1 and purk2 were not well separated (**Fig. 4B**), and the expression levels of *SORCS3* showed mosaic patterns across the 4 groups (cluster4, 6, 11 and 12, **Fig. 4E**). As additional controls, we performed clustering using the top 500 and 1000 Seurat HVGs. That did not result in any improvement (**S6 Fig**.).

**Fig 4.**
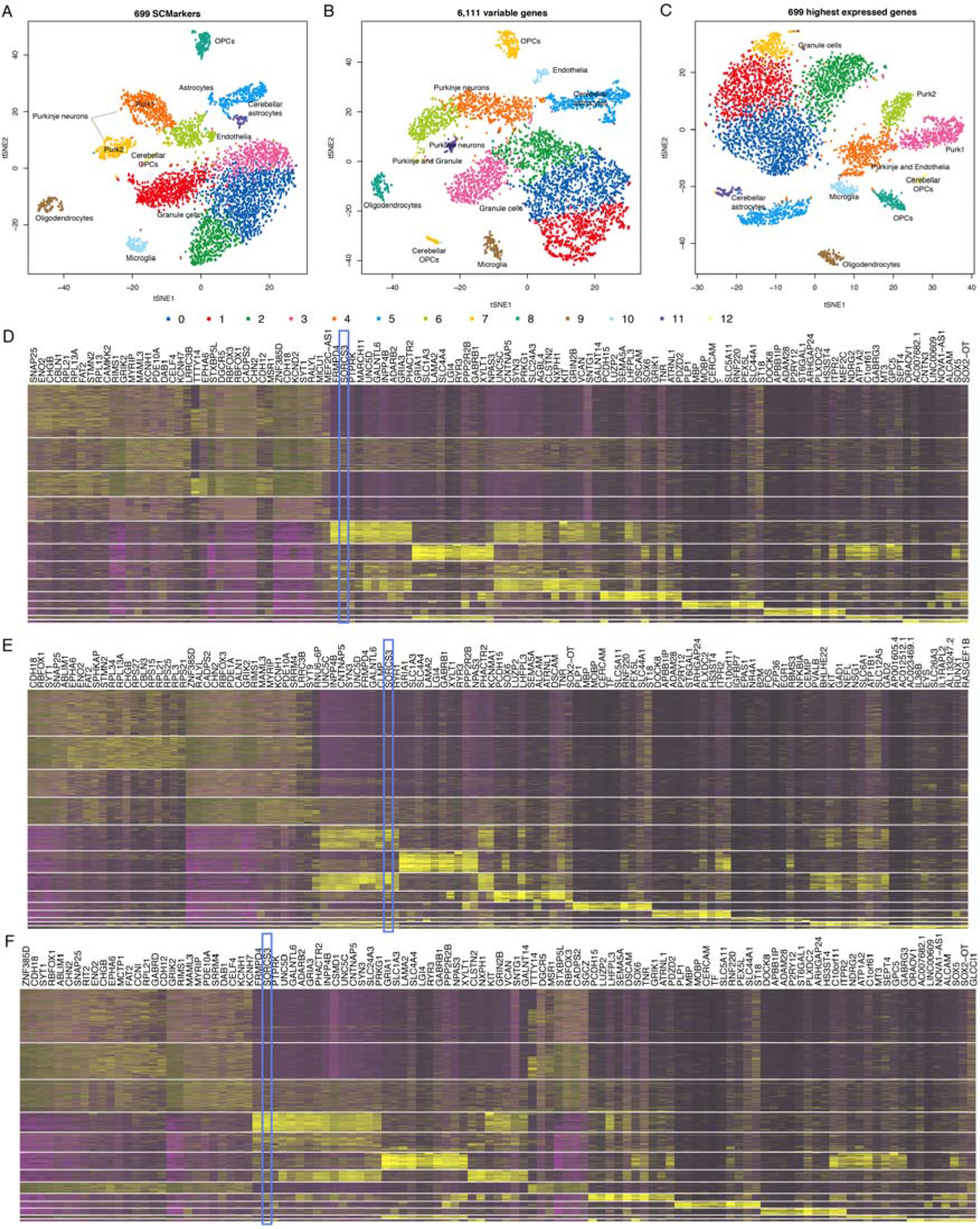
Results on the human brain data. Plotted in tSNE space are 5,602 cells in the cerebellar hemisphere of human brain tissue based on markers selected respectively by (**A**) SCMarker,(**B**) the highest variable genes and (**C**) the highest expressed genes, colored by performing clustering using Seurat. Cell types were labelled consistently as they were in the original paper. Also plotted are the heatmaps of the top 10 gene expression levels in each cluster derived respectively from (**D**) SCMarker,(**E**) the highest variable genes and (**F**) the highest expressed genes.

We then analysed the scRNA-seq data of 52,698 cells from 5 lung tumours generated by the 10X Chromium platform (10X Genomics) [35]. SCMarker identified 848 markers under the default parameters, while Seurat identified 1,832 HVGs. We also selected 848 HEGs for comparison. SCMarker led to 22 clearly distinguishable clusters, while the Seurat HVGs led to 12 and the HEGs led to 18 (**Fig. 5A** to **C**). The 10 highest expressed markers per cluster derived by SCMarker showed a high degree of cluster-specificity in the heatmap (**Fig. 5D**). Among the selected markers were the 16 known markers reported by the original study (Table 1). SCMarker also discovered multiple putative subtypes for some cell types, such as the T, B, fibroblast, and myeloid cells (Table 1, **Fig. 5A**). For example, cluster 7, 10, 13, 19, and 22 are the B cells expressing known surface marker *CD79A* (**Fig. 5A**), yet cells in cluster 7, 13, 19, and 22 are evidently different from cells in cluster 10, due to differential *IGHG1* and *BANK1* expression levels (**Fig. 5D**). For comparison, we selected the 10 highest expressed genes from the clusters determined by the Seurat HVGs and by the HEGs, respectively (**Fig. 5E** and **F**). They appeared non-specifically distributed across clusters (**Fig. 5E** and **F**). These genes also contained fewer known markers (Table 1). For example, cluster 3 determined by the Seurat HVGs contained markers (*CLDN5*, *CAV1* and *IFITM3*) from 3 cell-types (endothelia, alveolar and B cell, respectively). Most clusters expressed *IFITM3*, except for clusters 1 and 6 (**Fig. 5E**). Only T cell and fibroblast markers appeared to be cluster-specific. As additional controls, we also performed analysis using fewer (i.e., 500 and 1000) Seurat HVGs. That resulted in worse results with fewer known markers and marker-specific clusters (**S7 Fig**.).

**Fig. 5.**
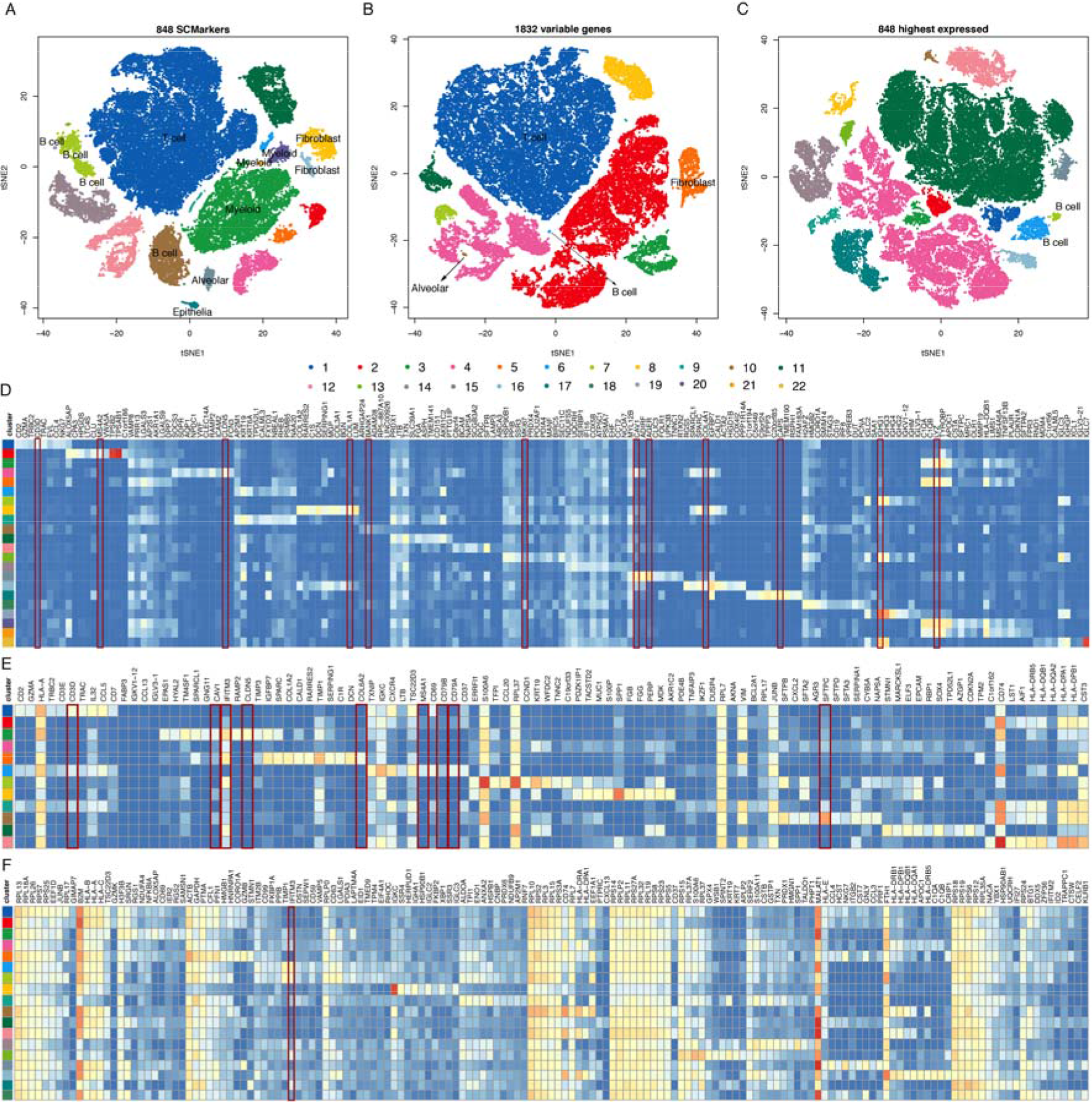
Results on the lung cancer data generated by Dropseq. Plotted in tSNE space are 52,698 cells of 6 different lung cancer patients, clustered based on markers selected respectively by (**A**) SCMarker, (**B**) the highest variable genes and (**C**) the highest expressed genes. Colors correspond to clusters determined by DBSCAN. Heatmaps of the average expression levels of the 10 highest expressed genes per cluster identified respectively by SCMarker (**D**), the highest variable genes (**E**) and the highest expressed genes (**F**). Cell types in (**A**) to (**C**) are labelled based on the known cell-type specific markers, which are highlighted in red boxes in (**D**) to (**F**).

**Table 1.** Known cell-type specific markers identified respectively by SCMarker and the highest variable gene approach.

| Known markers | Cell type | Cluster ID (SCMarker) | Cluster ID (highest variable) | Cluster ID (highest expressed) |
| --- | --- | --- | --- | --- |
| *CD3D* | T cell | 1 | 1 |  |
| *MKI67* | T cell | 18 |  |  |
| *IGHG1* | B cell | 7, 13, 19, 22 |  |  |

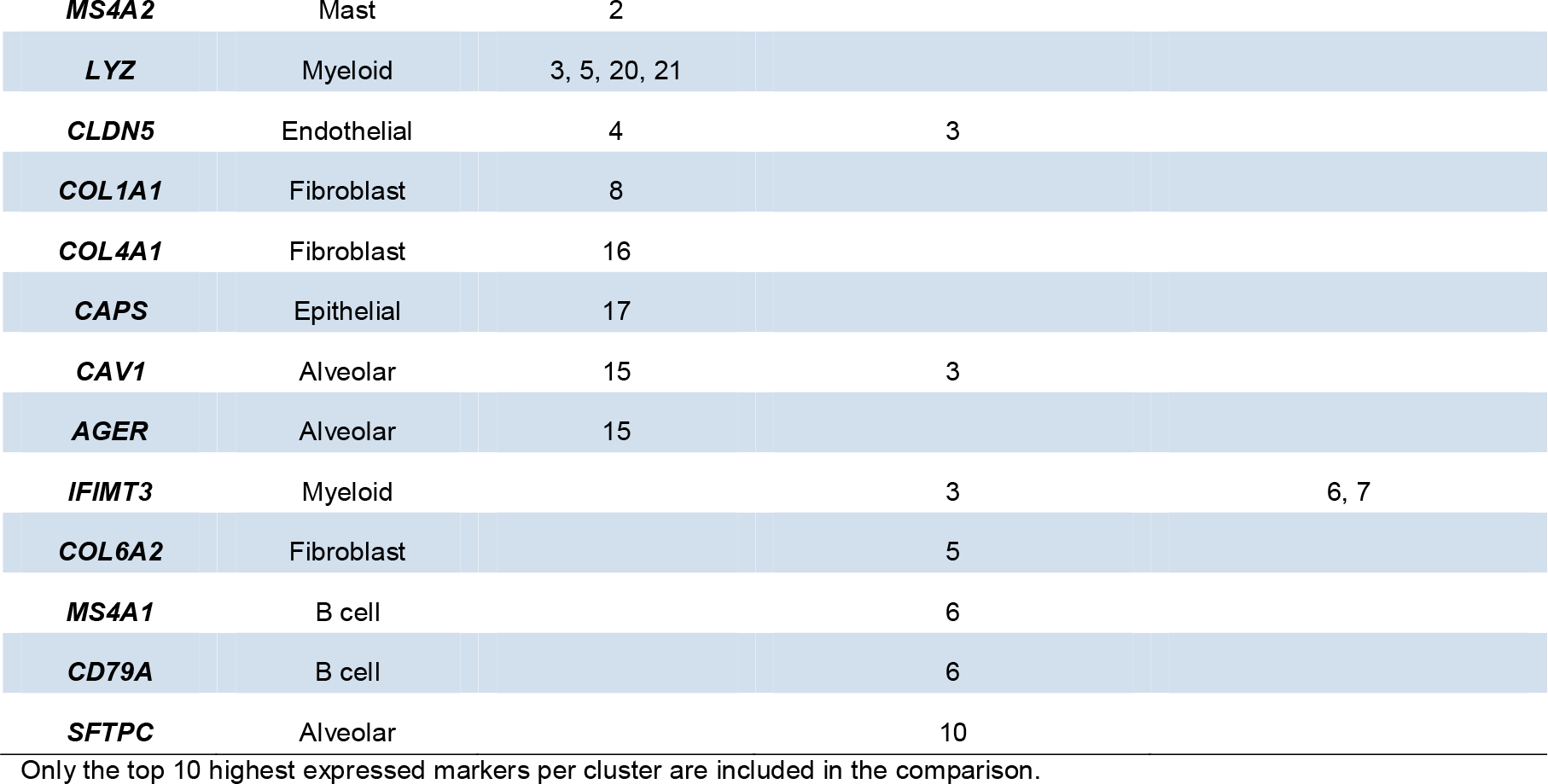

Overall, SCMarker demonstrated higher sensitivity and specificity for cell type and cell-type specific marker identification than the alternative approaches. Moreover, markers selected by SCMarker were more significant among genes which were identified by Seurat to define clusters (**S8 Fig.**).

## Discussion

In this manuscript, we reported a new bioinformatics tool, SCMarker, which performs *ab initio* cell-type discriminative marker selection from scRNA-seq data. SCMarker operates based on two new information-theoretic metrics: 1) bi/multi-modal distribution of subpopulation-discriminative gene expression in mixed cell populations and 2) co- or mutually-exclusively expressing gene pairs, which quantifies populational structural properties intrinsic to single-cell RNA-seq data. We found that SCMarker can consistently significantly boost cell-type identification accuracy in datasets from a variety of tissues such as cancer and brain, generated by both SMART-seq and Drop-seq platforms. Because SCMarker does not depend on any prior knowledge, we anticipate that it will prove most useful in discovery settings for analysing cell populations of a high degree of plasticity and heterogeneity [36]. SCMarker can potentially be expanded to analyse other types of single-cell data, including mass cytometry and single cell ATAC-seq (Assay for Transposase-Accessible Chromatin using sequencing) data [37,38]. It can be easily incorporated as a module into current scRNA-seq data analysis workflows to pre-process the cell-gene count/expression matrix before performing further downstream analysis.

## Acknowledgements

We thank Miao Qi and Yu-yu Lin for their important suggestions and feedback.

## Funding

This work was supported by a Chan-Zuckerberg Initiative award to Ken Chen.

## Supplementary Figure

**S1 Fig.** Distributions of expression levels of known marker genes (CD3G, CD8A, IL7R, MS4A1, CD19, CD79A, CD79B and PMEL) from melanoma data.

**S2 Fig.** Comparison of 3 marker selection methods for cell-type identification. Tested were a range of parameters and 5 clustering algorithms: k-means, Clara, hierarchical clustering (hc), DBSCAN, and Seurat. Plotted in heatmaps are the ARI values calculated based on markers selected respectively by SCMarker, the highest expressed genes and the highest variable genes from (A) the melanoma and (B) the head and neck cancer data. X and Y axes in the SCMarker panel indicate the *n* and *k* parameters used by SCMarker and the corresponding (equal number of markers) results in the highest expressed or the highest variable gene panels.

**S3 Fig.** Determining the optimal parameters. Plotted in the heatmaps are the number of selected markers for (**A**) the melanoma and (**B**) the head and neck cancer data over a range of *n* (X-axis) and *k* (Y-axis) parameters. Bars on the side and the top are the mean values in the corresponding rows and columns. Also plotted are clustering accuracy measured by the adjusted rand index (ARI), a metric that measures the similarity of two clustering results, for (**C**) the melanoma and (**D**) the head and neck cancer data over various *n* and *k* parameters.

**S4 Fig.** Validation of genes selected by SCMarker. Plotted in tSNE space are 5,902 cells from the head and neck cancer data, based on genes selected respectively by (**A**) SCMarker, (**B**) the highest expressed and (**C**) the highest variable genes.

**S5 Fig.** Gene set enrichment analysis (GSEA) of markers selected by 3 methods: SCMaker, the highest expressed and the highest variable genes from the (**a**) melanoma; and (**b**) the head and neck cancer data, respectively. Only the top 15 terms are shown. The darkness of the colors corresponds to −log10 P values.

**S6 Fig.** Results on the human brain tissue data. Plotted in tSNE space are 5,602 cells in the cerebellar hemisphere of human brain tissue based on the highest 500 (**A**) and 1000 (**B**) variable genes, colored by cell types from the original paper.

**S7 Fig.** Results on the lung cancer data. Plotted in tSNE space are 52,698 cells of 6 different lung cancer patients, clustered based on the highest 500 (**A**) and 1000 (**B**) variable genes. Colors correspond to clusters determined by DBSCAN. Heatmaps of the average expression levels of the 10 highest expressed genes per cluster identified respectively by the highest 500 (**C**) and 1000 (**D**) variable genes. Cell types in (A) and (B) are labelled based on the known cell-type specific markers, which are highlighted in red box in (C) and (D).

**S8 Fig.** The distribution of significance of markers identified by Seurat and overlaps with SCMarker in melanoma, head and neck cancer (HNSCC), brain tissue and lung cancer.

