## Supplementary Figures and Tables for "SCMarker: ab initio marker selection for single cell transcriptome profiling"

**S1 Table. Clustering methods.**

| **Method** | **Description** |
| --- | --- |
| Ascend | PCA dimension reduction and iterative hierarchical cluster |
| CIDR | PCA dimension reduction and hierarchical clustering |
| FlowSOM | PCA dimension reduction, then self-organizing maps and hierarchical consensus meta-clustering |
| PCAHC | PCA dimension reduction and hierarchical clustering |
| PCAKmeans | PCA dimension reduction and Kmeans |
| pcaReduce | PCA dimension reduction and iterative Kmeans |
| RtsneKmeans | t-SNE dimension reduction and Kmeans |
| SAFE | Ensemble clustering using SC3, Seurat, and tSNE+Kmeans |
| SC3 | PCA dimension reduction, then kmeans and different dimensions and hierarchical clustering on consensus |
| SC3svm | SC3 and SVM |
| Seurat | PCA dimension reduction and nearest neighbor graph clustering |
| TSCAN | PCA dimension reduction and model based clustering |

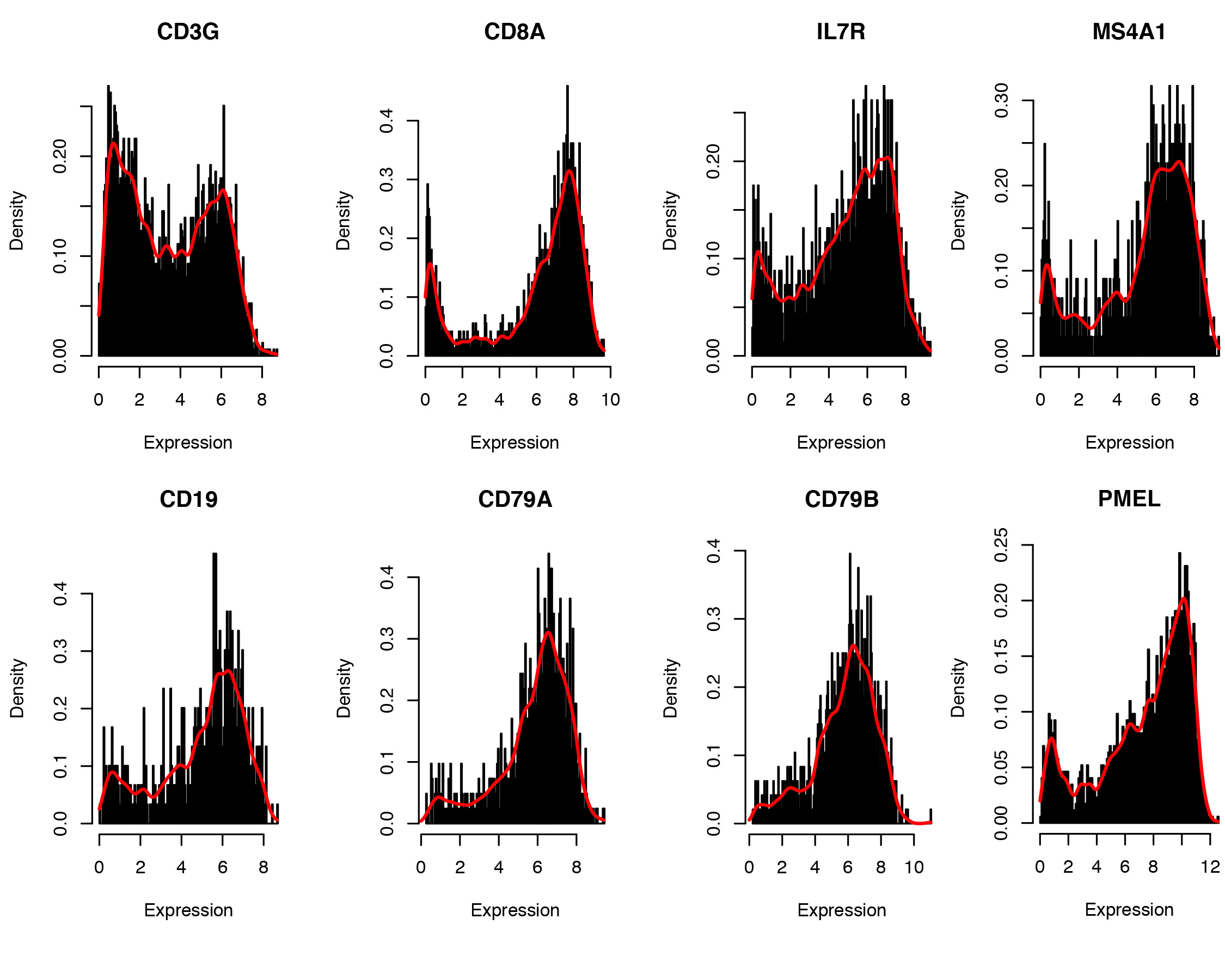

**S1 Fig. The** expression levels of most known marker genes (*CD3G, CD8A, IL7R, MS4A1, CD19, CD79A, CD79B* and *PMEL*) follow a bi/multi-modal distribution in the melanoma data.

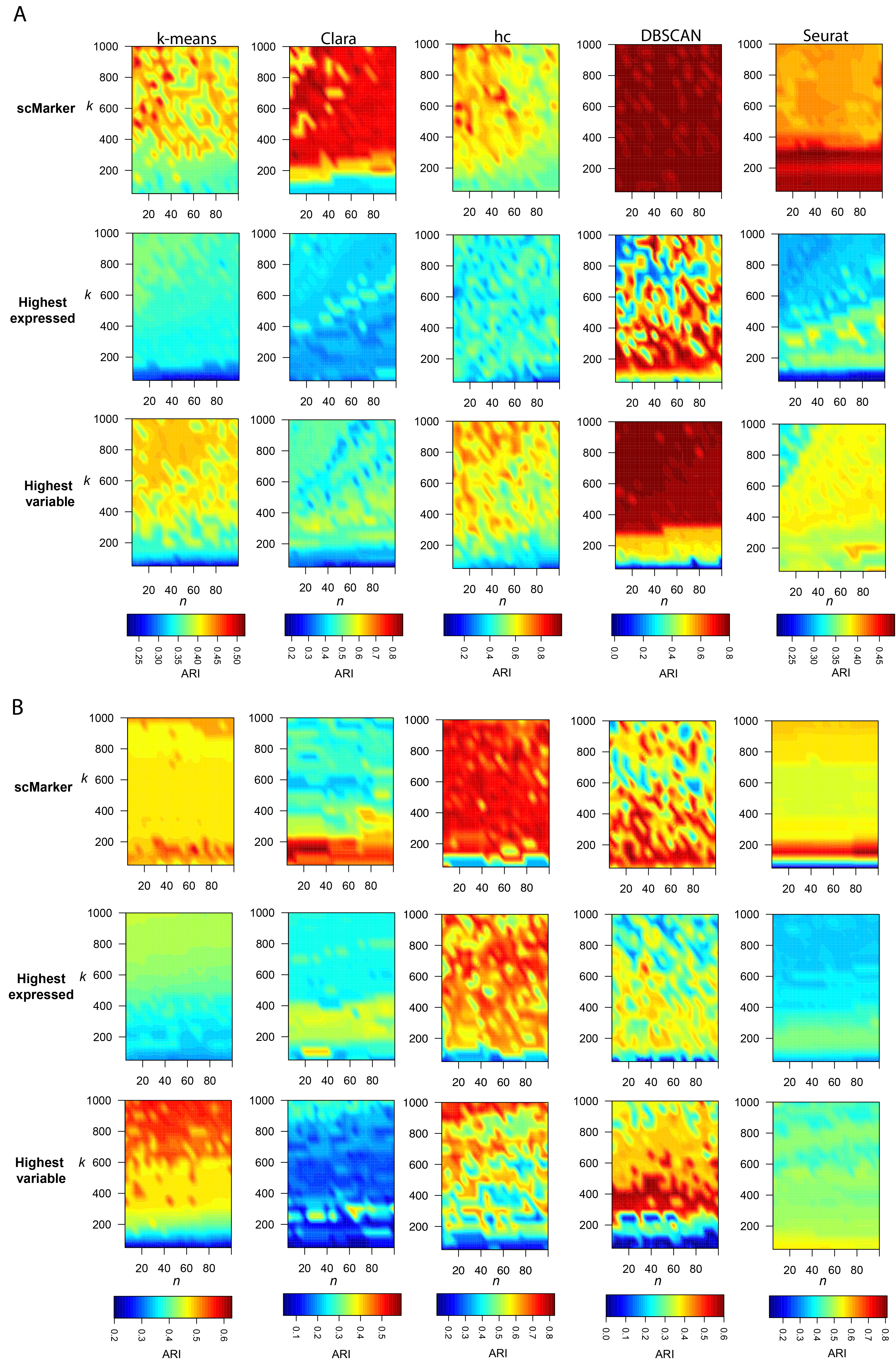

**S2 Fig.** Comparison of 3 marker selection methods for cell-type identification. Tested were a range of parameters and 5 clustering algorithms: k-means, Clara, hierarchical clustering (hc), DBSCAN, and Seurat. Plotted in heatmaps are the ARI values calculated based on markers selected respectively by SCMarker, the highest expressed genes and the highest variable genes from (A) the melanoma and (B) the head and neck cancer data. X and Y axes in the SCMarker panel indicate the $n$ and $k$ parameters used by SCMarker and the corresponding (equal number of markers) results in the highest expressed or the highest variable gene panels.

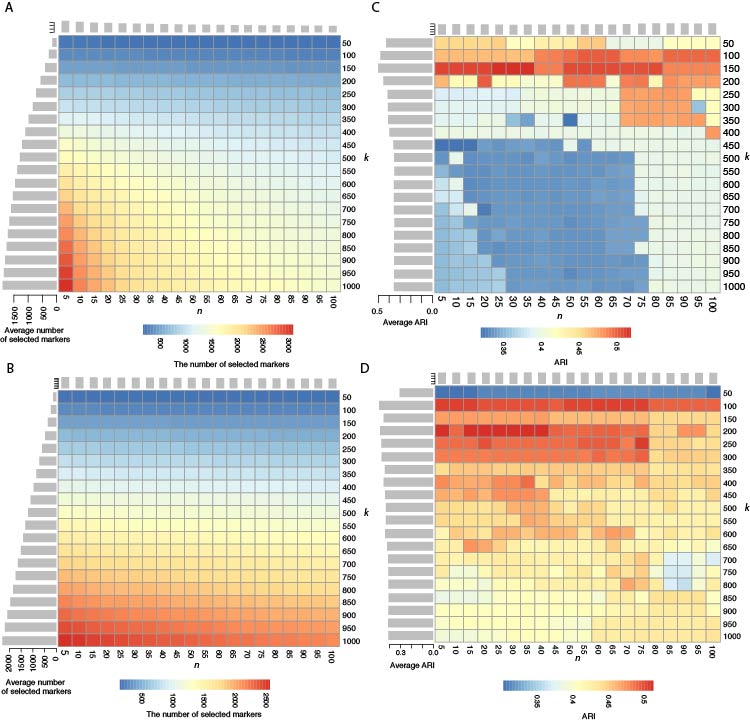

**S3 Fig.** Determining the optimal parameters. Plotted in the heatmaps are the number of selected markers for (**A**) the melanoma and (**B**) the head and neck cancer data over a range of $n$ (X-axis) and $k$ (Y-axis) parameters. Bars on the side and the top are the mean values in the corresponding rows and columns. Also plotted are clustering accuracy measured by the adjusted rand index (ARI), a metric that measures the similarity of two clustering results, for (**C**) the melanoma and (**D**) the head and neck cancer data over various $n$ and $k$ parameters.

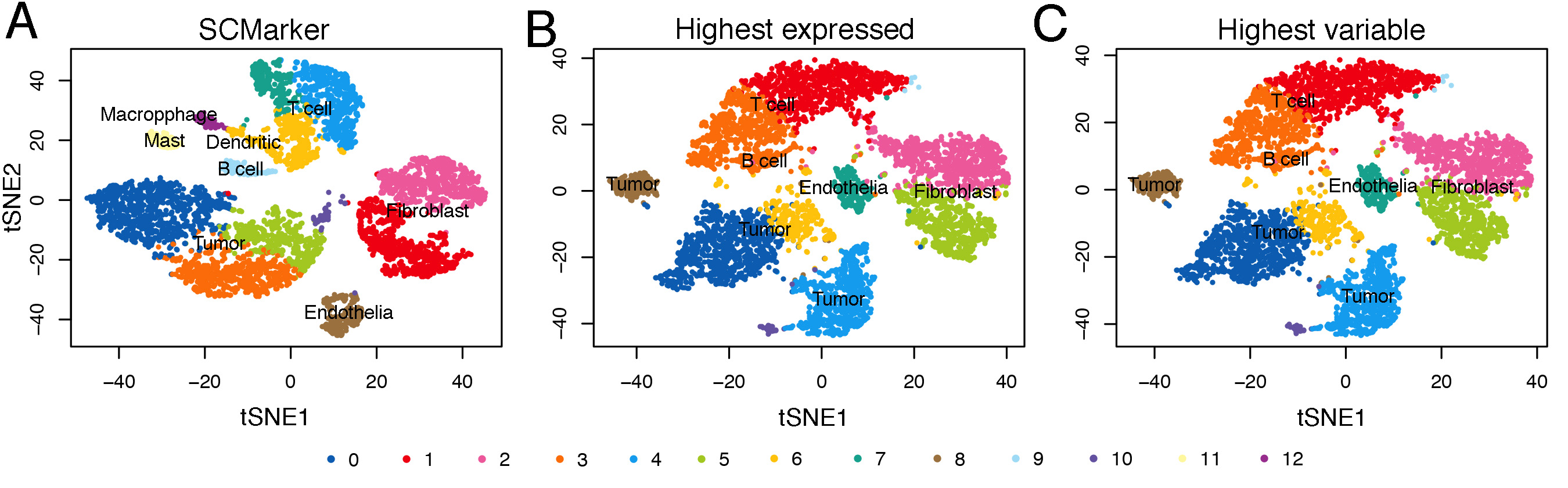

**S4 Fig.** Validation of genes selected by SCMarker. Plotted in tSNE space are 5,902 cells from the head and neck cancer data, based on genes selected respectively by (**A**) SCMarker, (**B**) the highest expressed and (**C**) the highest variable genes.

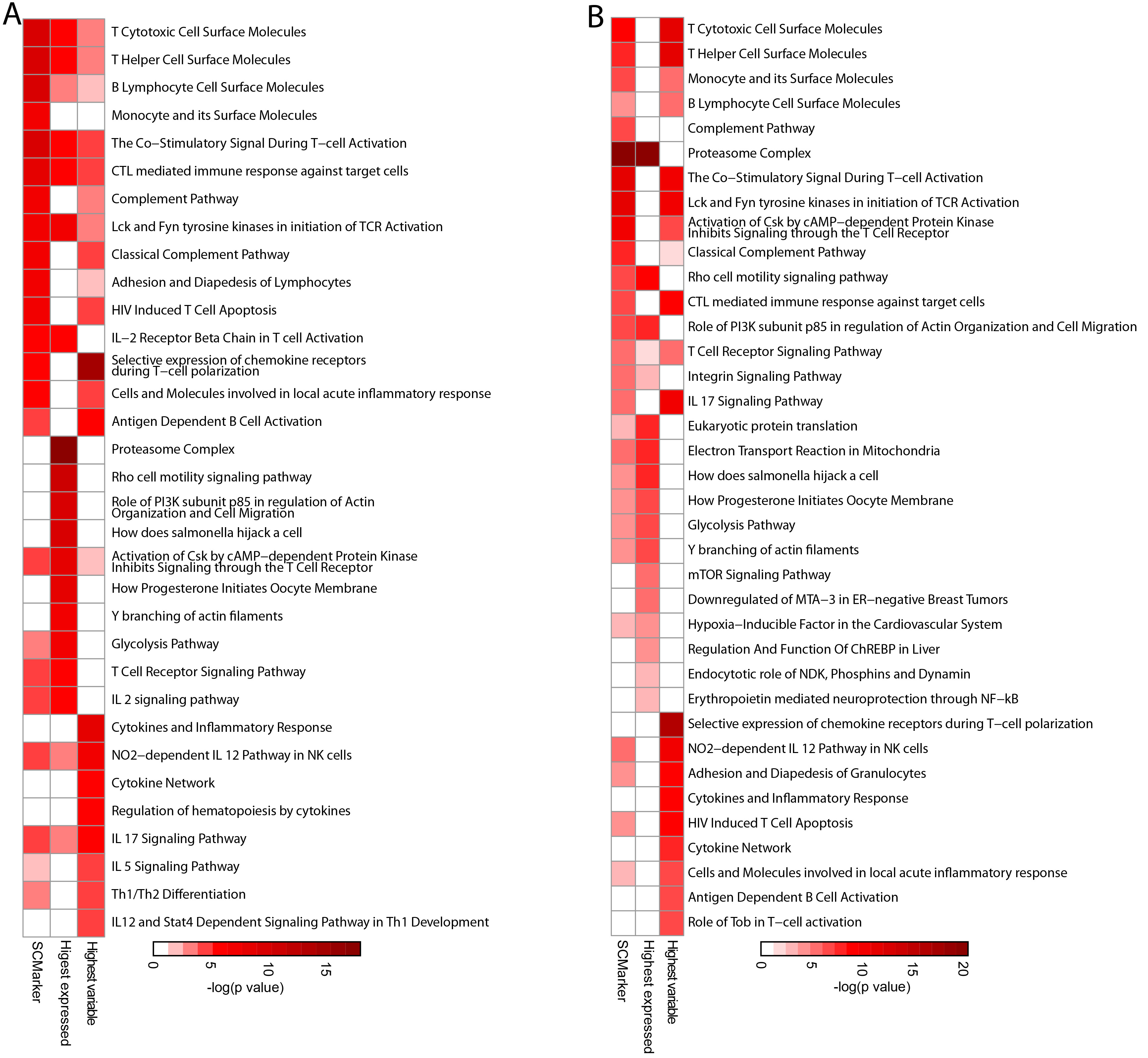

**S5 Fig.** Gene set enrichment analysis (GSEA) of markers selected by 3 methods: SCMaker, the highest expressed and the highest variable genes from the (**A**) melanoma; and (**B**) the head and neck cancer data, respectively. Only the top 15 terms are shown. The darkness of the colors corresponds to -log10 P values.

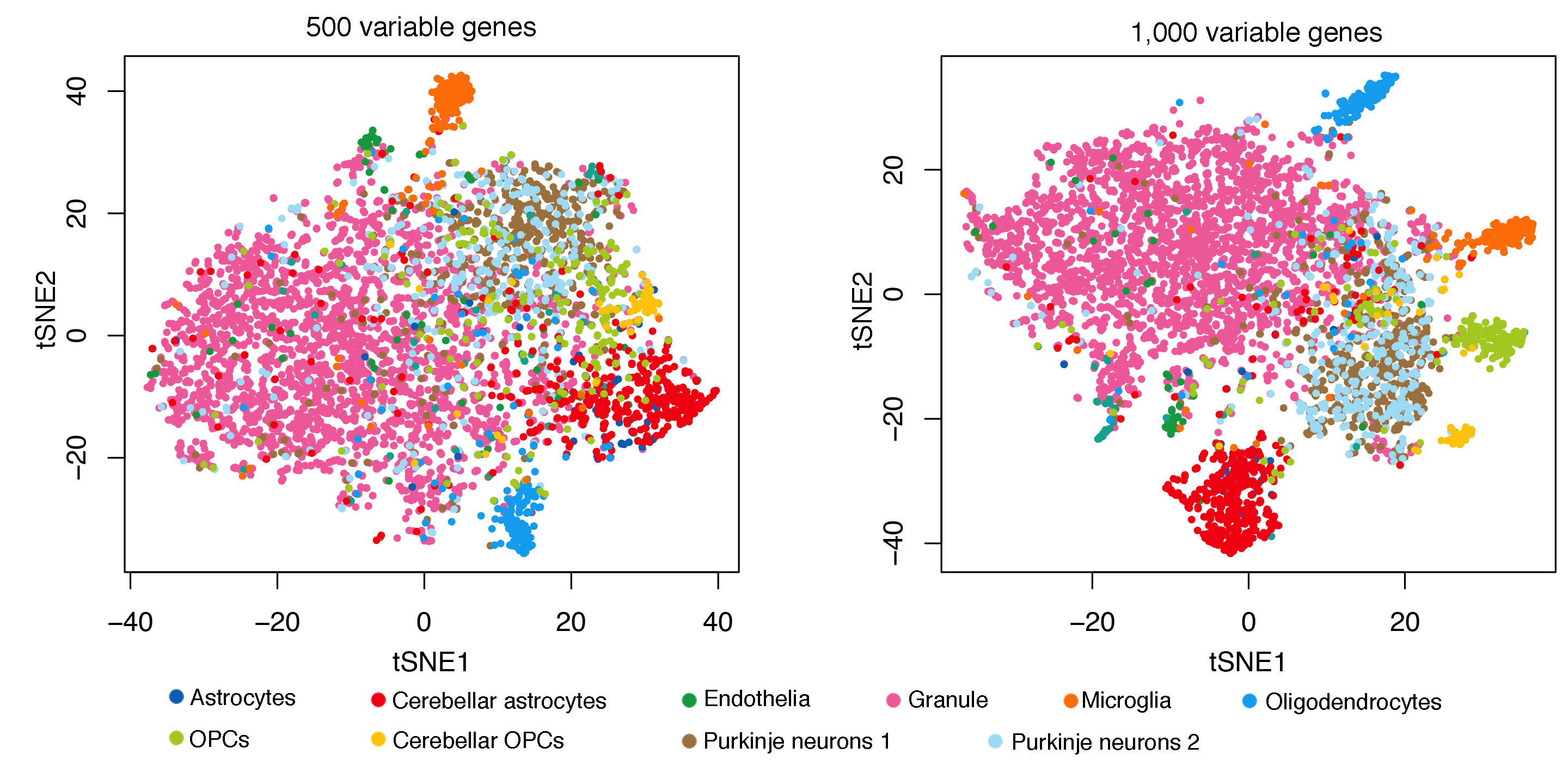

**S6 Fig.** Results on the human brain tissue data. Plotted in tSNE space are 5,602 cells in the cerebellar hemisphere of human brain tissue based on the highest 500 (**A**) and 1000 (**B**) variable genes, colored by cell types from the original paper.

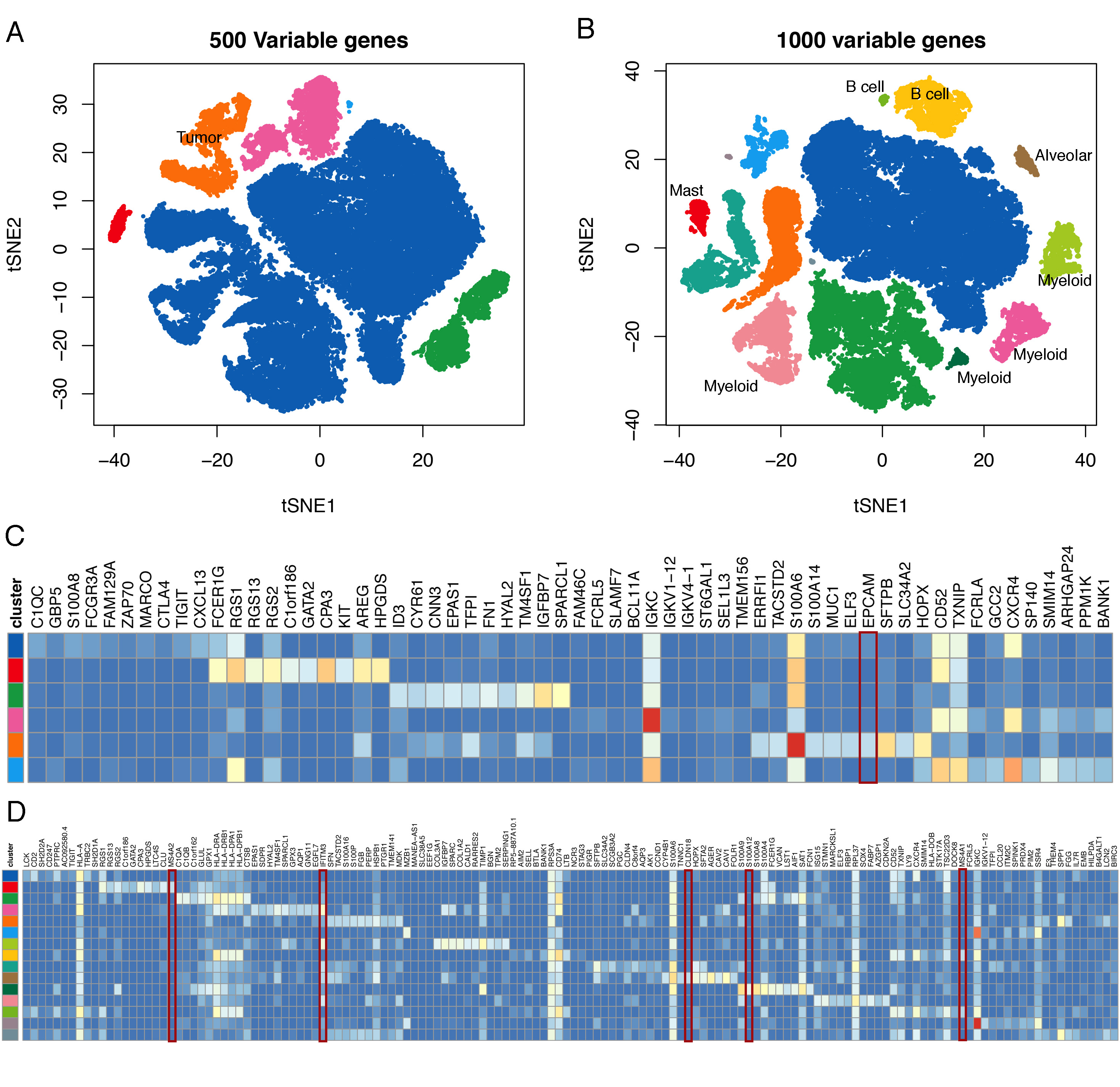

**S7 Fig.** Results on the lung cancer data. Plotted in tSNE space are 52,698 cells of 6 different lung cancer patients, clustered based on the highest 500 (**A**) and 1000 (**B**) variable genes. Colors correspond to clusters determined by DBSCAN. Heatmaps of the average expression levels of the 10 highest expressed genes per cluster identified respectively by the highest 500 (**C**) and 1000 (**D**) variable genes. Cell types in (A) and (B) are labelled based on the known cell-type specific markers, which are highlighted in red box in (C) and (D).

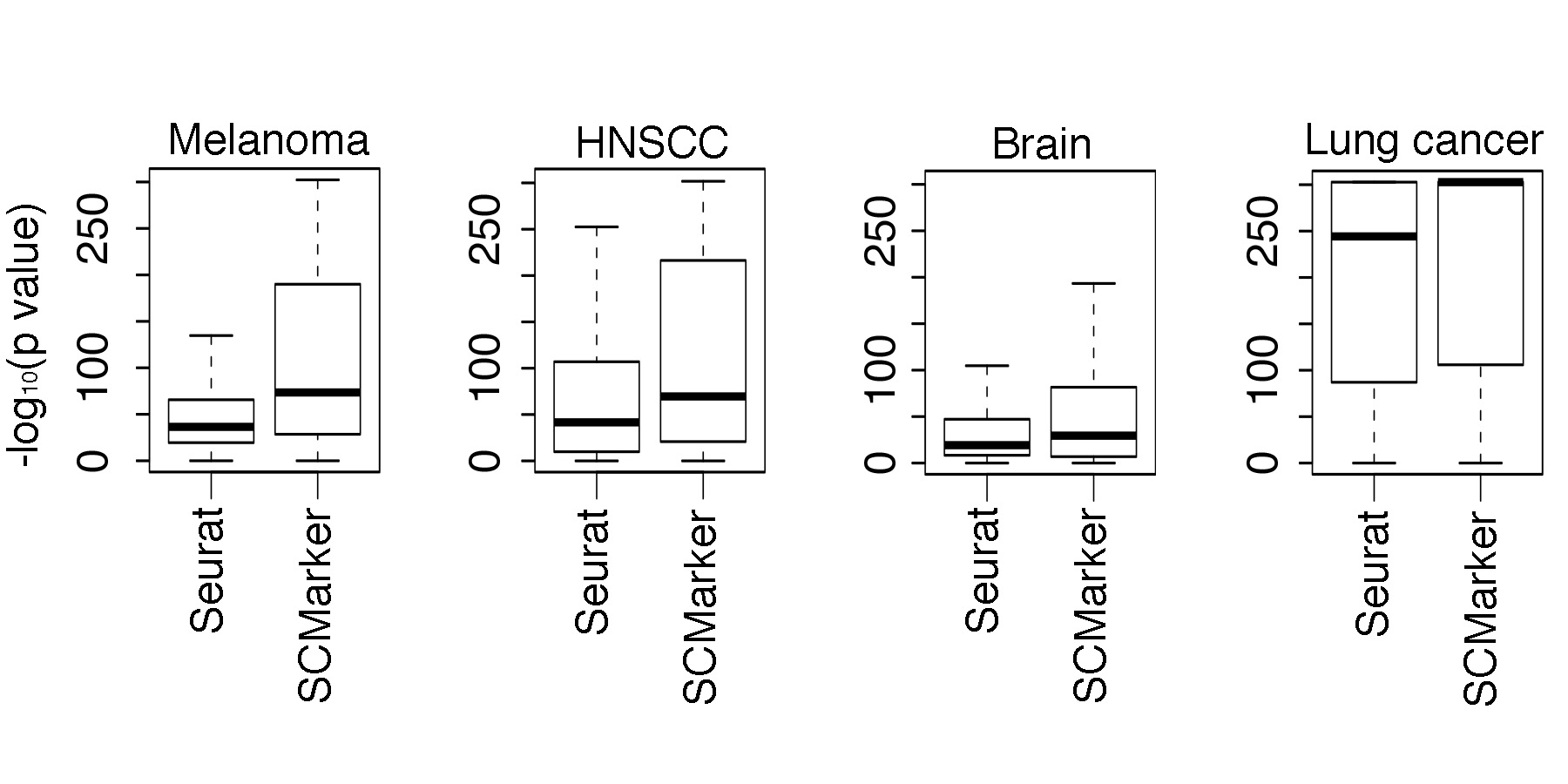

**S8 Fig.** The distribution of sSignificance of overlaps between markers identified by Seurat and SCMarker overlaps with SCMarkerand the known markers in the melanoma, head and neck cancer (HNSCC), brain tissue and lung cancer data.
